# A unified sequence catalogue of over 280,000 genomes obtained from the human gut microbiome

**DOI:** 10.1101/762682

**Authors:** Alexandre Almeida, Stephen Nayfach, Miguel Boland, Francesco Strozzi, Martin Beracochea, Zhou Jason Shi, Katherine S. Pollard, Donovan H. Parks, Philip Hugenholtz, Nicola Segata, Nikos C. Kyrpides, Robert D. Finn

## Abstract

Comprehensive reference data is essential for accurate taxonomic and functional characterization of the human gut microbiome. Here we present the Unified Human Gastrointestinal Genome (UHGG) collection, a resource combining 286,997 genomes representing 4,644 prokaryotic species from the human gut. These genomes contain over 625 million protein sequences used to generate the Unified Human Gastrointestinal Protein (UHGP) catalogue, a collection that more than doubles the number of gut protein clusters over the Integrated Gene Catalogue. We find that a large portion of the human gut microbiome remains to be fully explored, with over 70% of the UHGG species lacking cultured representatives, and 40% of the UHGP missing meaningful functional annotations. Intra-species genomic variation analyses revealed a large reservoir of accessory genes and single-nucleotide variants, many of which were specific to individual human populations. These freely available genomic resources should greatly facilitate investigations into the human gut microbiome.

## Main

The human gut microbiome has been implicated in important phenotypes related to human health and disease^1,2^. However, incomplete reference data that are missing microbial diversity^3^ hamper our understanding of the roles of individual microbiome species, their interactions and functions. Hence, establishing a comprehensive collection of microbial reference genomes and genes is an important step for accurate characterization of the taxonomic and functional repertoire of the intestinal microbial ecosystem.

The Human Microbiome Project (HMP)^4^ was a pioneering initiative to enrich our knowledge of human-associated microbiota diversity. Hundreds of genomes from bacterial species with no sequenced representatives were obtained as part of this project, allowing their use for the first time in reference-based metagenomic studies. The Integrated Gene Catalogue (IGC)^5^ was subsequently created, combining the sequence data available from the HMP and the Metagenomics of the Human Intestinal Tract (MetaHIT)^6^ consortium. This gene catalogue has been applied successfully to the study of microbiome associations in different clinical contexts^7^, revealing microbial composition signatures linked to type 2 diabetes^8^, obesity^9^ and other diseases^10^. But, as the IGC comprises genes with no direct link to their genome of origin, it lacks contextual data to perform a high-resolution taxonomic classification, establish genetic linkage and deduce complete functional pathways on a genomic basis.

Culturing studies have continued to unveil new insights into the biology of our gut communities^11,12^ and are essential for applications in research and biotechnology. However, the advent of high-throughput sequencing and new metagenomic analysis methods — namely involving genome assembly and binning — has transformed our understanding of the microbiome composition both in humans and other environments^13–15^. Metagenomic analyses are able to capture substantial microbial diversity not easily accessible by cultivation by directly analysing the sample genetic material without the need for culturing, though biases do exist^16^. Recent studies have massively expanded the known species repertoire of the human gut, making available unprecedented numbers of new cultured and uncultured genomes^16–20^. Two culturing efforts isolated and sequenced over 500 human gut-associated bacterial genomes each^18,20^, while three independent studies^16,17,19^ reconstructed 60,000–150,000 microbial metagenome-assembled genomes (MAGs) from public human microbiome data, most of which belong to species lacking cultured representatives. Combining these individual efforts and establishing a unified non-redundant dataset of human gut genomes is essential for driving future microbiome studies. To accomplish this, we compiled and analysed 286,997 genomes and 625,251,941 genes from human gut microbiome datasets to generate the Unified Human Gastrointestinal Genome (UHGG) and Protein (UHGP) catalogues, the most comprehensive sequence resources of the human gut microbiome established to date.

## Results

### The UHGG represents over 280,000 human gut microbial genomes

We first gathered all prokaryotic isolate genomes and MAGs from the human gut microbiome (publicly available as of March 2019). We compiled the isolate genomes from the Human Gastrointestinal Bacteria Culture Collection (HBC)^18^, the Culturable Genome Reference (CGR)^20^, as well as cultured human gut genomes available in the NCBI^21^, PATRIC^22^ and IMG^23^ repositories which include genomes from several other large studies^11,12,24^. In addition, we included all of the gut MAGs generated in Pasolli, et al.^19^ (CIBIO), Almeida, et al.^17^ (EBI) and Nayfach, et al.^16^ (HGM). To standardize the genome quality across all sets, we used thresholds of >50% genome completeness and <5% contamination, combined with an estimated quality score (completeness – 5 × contamination) >50. Final numbers of genomes matching these criteria were: 734 (HBC), 1,519 (CGR), 651 (NCBI), 7,744 (PATRIC/IMG), 137,474 (CIBIO), 87,386 (EBI) and 51,489 (HGM), resulting in a total of 286,997 genome sequences (Fig. 1a and Supplementary Table 1). Genomes were recovered in samples from a total of 31 countries across six continents (Africa, Asia, Europe, North America, South America and Oceania), but the majority originated from samples collected in China, Denmark, Spain and the United States (Fig. 1b).

**Figure 1.**
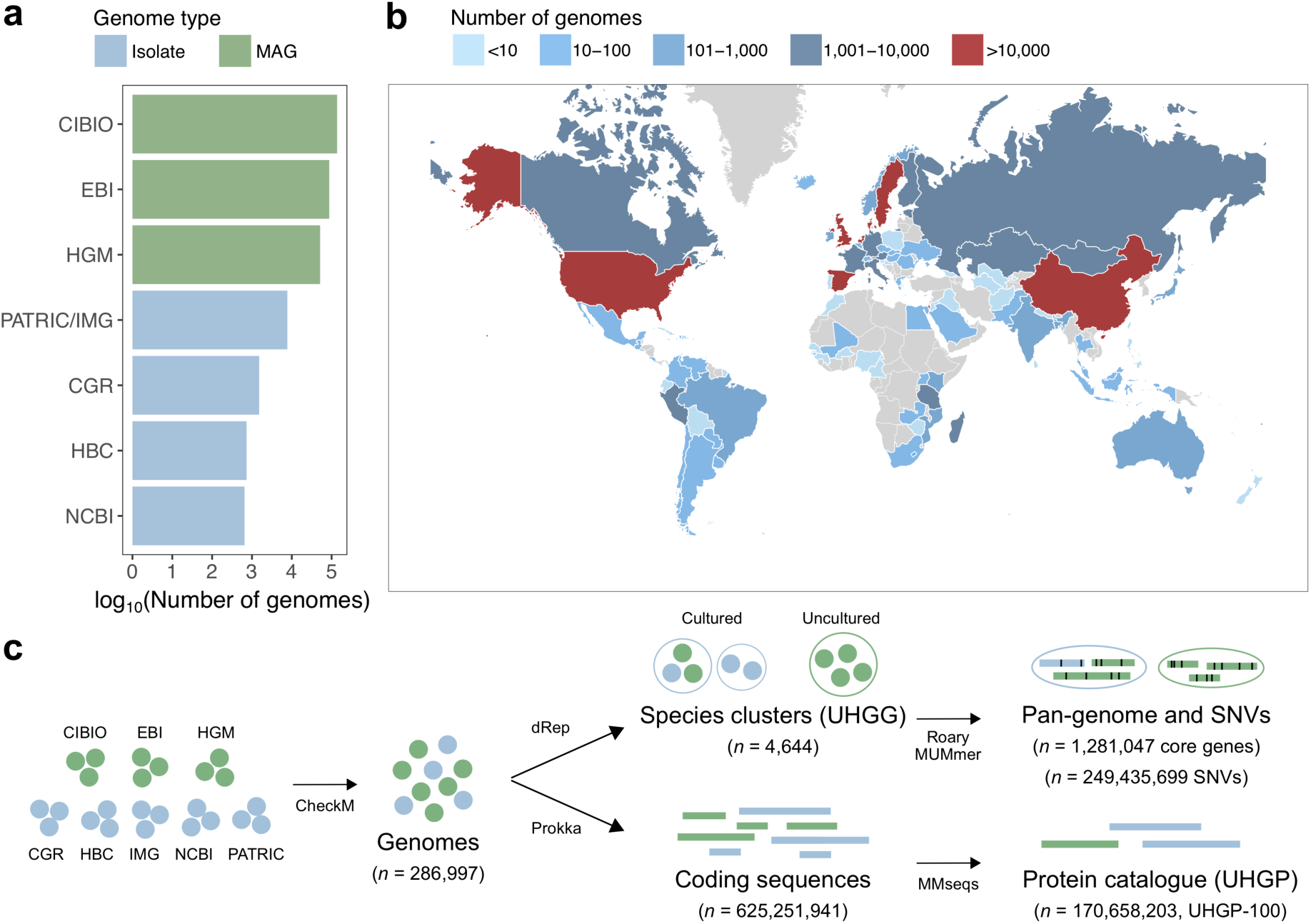
The unified sequence catalogue of the human gut microbiome. **a**, Number of gut genomes per each study set used to generate the sequence catalogues, coloured according to whether they represent isolate genomes or metagenome-assembled genomes (MAGs). **b**, Geographic distribution of the number of genomes retrieved per country. **c**, Overview of the methods used to generate the genome (UHGG) and protein sequence (UHGP) catalogue. Genomes retrieved from public datasets were first quality-controlled by CheckM. Filtered genomes were clustered at an estimated species-level (95% average nucleotide identity) and their intra-species diversity was assessed (genes from conspecific genomes were clustered at a 90% protein identity). In parallel, a non-redundant protein catalogue was generated from all the coding sequences of the 286,997 genomes at 100% (UHGP-100, *n* = 170,658,203), 95% (UHGP-95, *n* = 20,240,320), 90% (UHGP-90, *n* = 13,910,025) and 50% (UHGP-50, *n* = 4,736,012) protein identity.

To determine how many species were included in this gut reference collection, we clustered all 286,997 genomes using a multi-step distance-based approach (see ‘Methods’) with an average nucleotide identity (ANI) threshold of 95% over at least a 30% alignment fraction^25^. The clustering procedure resulted in a total of 4,644 inferred prokaryotic species (4,616 bacterial and 28 archaeal, Supplementary Table 2). We found the species clustering results to be highly consistent with those previously obtained^16,17,19^ (Supplementary Table 3). The best quality genome from each species cluster was selected as its representative on the basis of genome completeness, contamination and assembly N50 (with isolate genomes always given preference over MAGs), and the final set was used to generate the Unified Human Gastrointestinal Genome (UHGG) catalogue (Fig. 1c). Out of the 4,644 species-level genomes, 3,207 were >90% complete (interquartile range, IQR = 87.2–98.8%) and <5% contaminated (IQR = 0.0–1.34%), with 573 of those having the 5S, 16S and 23S rRNA genes together with at least 18 of the standard tRNAs (Supplementary Fig. 1). These 573 genomes satisfy the “high quality” criteria set for MAGs by the Genomic Standards Consortium^26^. Thereafter, we classified each species representative using the Genome Taxonomy Database^27^ Toolkit (Supplementary Fig. 2), a standardized taxonomic framework based on a concatenated protein phylogeny representing >140,000 public prokaryote genomes, fully resolved to the species level (see ‘Methods’ for details on the taxonomy nomenclature used). However, over 60% of the gut genomes could not be assigned to an existing species, confirming the majority of the UHGG species lack representation in current reference databases.

### Comparison of species recovered in individual studies

We investigated how many of the 4,644 gut species were found in the different study collections in order to determine their level of overlap and reproducibility, as well as the ratio between cultured and uncultured species (Fig. 2a). The largest intersection was between the collections of MAGs, with the same 1,081 species detected independently in the CIBIO, EBI and HGM datasets, but not in any of the cultured genome studies. By restricting the analysis to genomes recovered from 1,554 samples common to all three MAG studies, we found that 93-97% of species from each set were detected in at least one other MAG collection, and 79-86% across all three (Supplementary Fig. 3a). Similar level of species overlap was observed when comparing studies on a per-sample basis (Supplementary Fig. 3b). Further, conspecific genomes recovered from the same samples across different studies shared a median ANI and aligned fraction of 99.9% and 92.1%, respectively (Supplementary Fig. 3c). These results emphasize the reproducibility of the different assembly and binning methods used in the large-scale studies of human gut MAGs^16,17,19^. Importantly, rarefaction analysis indicates the number of uncultured species detected has not reached a saturation point, meaning additional species remain to be discovered (Fig. 2b). However, these most likely represent rarer members of the human gut microbiome, as the number of species is closer to saturating when only considering those with at least two conspecific genomes.

**Figure 2.**
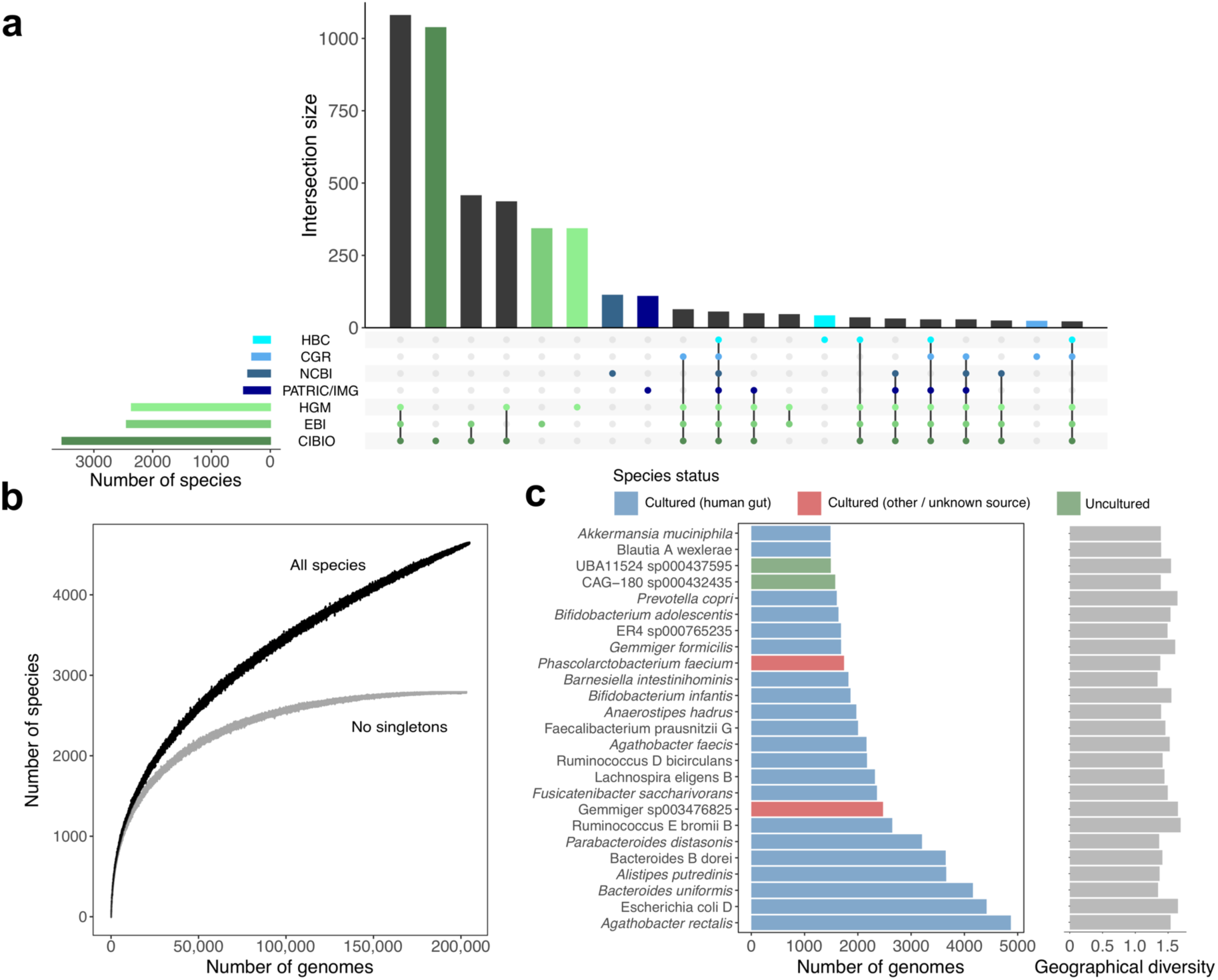
Intersection and frequency of species across studies. **a**, Number of species found across the genome study sets here used, ordered by their level of overlap. Vertical bars represent the number of species shared between the study sets highlighted in the lower panel. **b**, Rarefaction curve of the number of species detected as a function of the number of non-redundant genomes analysed. Curves are depicted both for all the UHGG species, and after excluding singleton species (represented by only one genome). **c**, Number of non-redundant genomes detected per species (left) alongside the degree of geographical diversity (calculated with the Shannon diversity index, right).

We also investigated the intersection between the three large culture-based datasets: the HBC, CGR and the NCBI (which contains gut genomes from the Human Microbiome Project, HMP^4^). Unlike the MAGs, the majority of cultured species were unique within a single collection (486/698; 70%), with only 70 (10%) being common to all three collections (Supplementary Fig. 3d). This may be due to varied geographical sampling between the collections (Asia, Europe and North America) or highlight the stochastic nature of culture-based studies.

By calculating the number of genomes contained within each cultured and uncultured species, we found that species containing isolate genomes represented the largest clusters, while those exclusively encompassing MAGs tended to be the rarest, as discussed previously^16,17,19^. For example, only two of the 25 largest bacterial clusters were exclusively represented by MAGs (Fig. 2c), with 1,212 uncultured species represented by a single genome (80% of which originated from samples only analysed in one of the MAG studies; Supplementary Fig. 4). The bacterial species most represented in our collection were *Agathobacter rectalis* (recently reclassified from *Eubacterium rectale*^28^), Escherichia coli D and *Bacteroides uniformis* (Fig. 2c, Supplementary Fig. 5 and Supplementary Table 2), whereas the most frequently recovered archaeal species was Methanobrevibacter A smithii, with 608 genomes found across all six continents (Supplementary Fig. 6). The largest species clusters displayed similarly high levels of geographical distribution, indicating the most highly represented species were not restricted to individual locations (Fig. 2c and Supplementary Fig. 5b).

### Most gut microbial species still lack isolate genomes

We found that 3,750 (81%) of the species in the UHGG did not have a representative in any of the human gut culture databases. To extend the search to isolate genomes from other environments or lacking information on their isolation source, we compared the UHGG catalogue to all NCBI RefSeq isolate genomes. We identified an additional set of 438 species closely matching cultured genomes, leaving 3,312 (71%) of UHGG species as uncultured (Supplementary Table 2).

The phylogenetic distribution of the 4,616 bacterial (Fig. 3a) and the 28 archaeal species (Supplementary Fig. 6) revealed that uncultured species exclusively represented 66% and 31% of the phylogenetic diversity of Bacteria and Archaea, respectively, with several phyla lacking cultured representatives (Fig. 3b). The four largest monophyletic groups lacking cultured genomes were the 4C28d-15 order (167 species, recently proposed as the novel order Comantemales ord. nov.^29^; Fig. 3c), order RF39 (139 species), family CAG-272 (88 species), and order Gastranaerophilales (67 species). While none have been successfully cultured, several have been described in the literature, including RF39^16^ and Gastranaerophilales (previously classified as a lineage in the Melainabacteria^30^) which are characterized by highly reduced genomes with numerous auxotrophies. This analysis suggests that, despite recent culture-based studies^11,12,18,20^, much of the diversity in the gut microbiome remains uncultured, including several large and prevalent clades.

**Figure 3.**
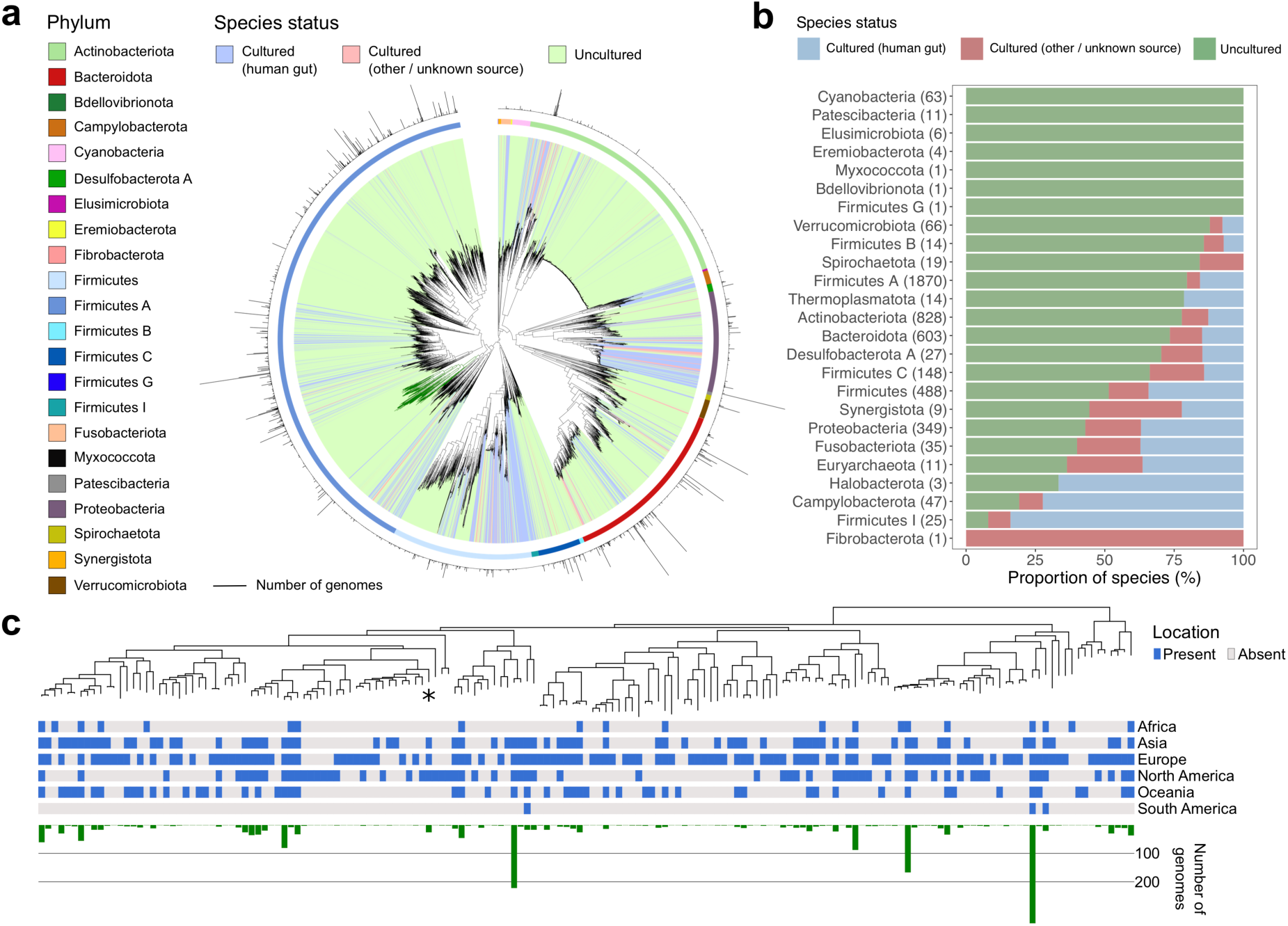
Uncultured species are predominant among human gut phyla. **a**, Maximum-likelihood phylogenetic tree of the 4,616 bacterial species detected in the human gut. Clades are coloured by species cultured status with outer circles depicting the GTDB phylum annotation. Bar graphs in the outermost layer indicate the number of genomes from each species. The order Comantemales ord. nov. is highlighted with dark green branches. **b**, Proportion of species within the 25 prokaryotic phyla detected according to their cultured status. Numbers in brackets represent the total number of species in the corresponding phylum. **c**, Phylogenetic tree of species belonging to the order Comantemales ord. nov. (phylum Firmicutes A), the largest phylogenetic group exclusively represented by uncultured species. The geographic distribution of each species and the number of genomes recovered is represented below the tree. The species previously classified as *Candidatus* Borkfalki ceftriaxensis is indicated with an asterisk.

### The UHGP expands the protein universe in the human gut microbiome

Metagenomic approaches have the ability to leverage gene content information not only for more precise taxonomic analysis, but to also predict the functional capacity of individual species of interest compared to marker gene-based methods (e.g. relying solely on the 16S rRNA gene or a limited number of diagnostic genes). We built the Unified Human Gastrointestinal Protein (UHGP) catalogue with a total of 625,251,941 full-length protein sequences predicted from the 286,997 genomes here analysed. These were clustered at 50% (UHGP-50), 90% (UHGP-90), 95% (UHGP-90) and 100% (UHGP-100) amino acid identity, generating between 5 to 171 million protein clusters (Fig. 1c and Fig. 4a).

**Figure 4.**
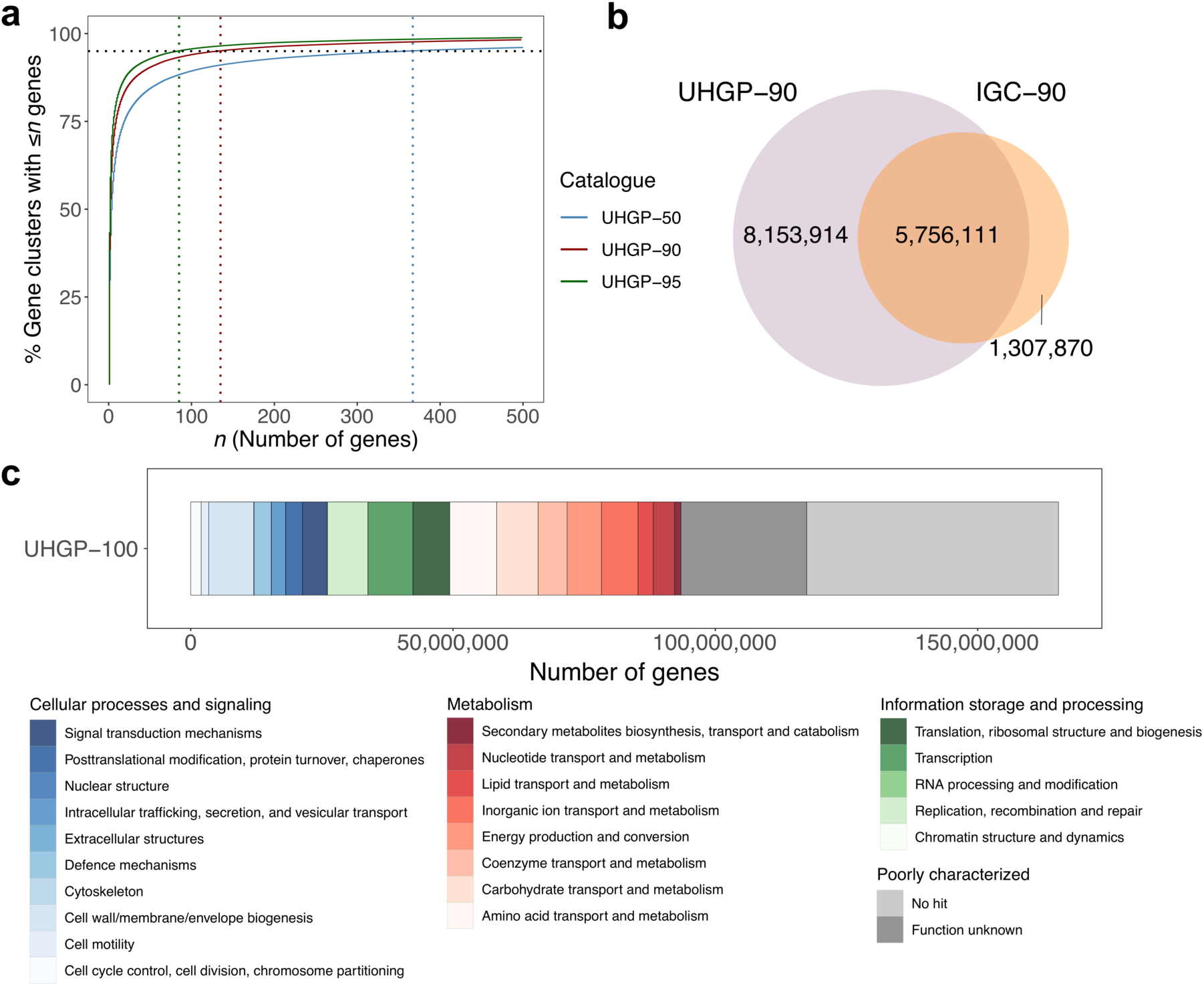
The UHGP improves coverage of the human gut protein landscape. **a**, Cumulative distribution curve of the number and size of the gene clusters of the UHGP-95 (*n* = 20,240,320), UHGP-90 (*n* = 13,910,025) and UHGP-50 (*n* = 4,736,012). Dashed vertical lines indicate the cluster size below which 90% of the gene clusters can be found. **b**, Overlap between the UHGP (purple) and IGC (orange), both clustered at 90% amino acid identity. **c**, COG functional annotation results of the unified gastrointestinal protein catalogue clustered at 100% amino acid identity (UHGP-100).

To determine how comprehensive the UHGP was when compared to existing human gut gene catalogues, we combined the UHGP-90 (*n* = 13,910,025 protein clusters) together with the Integrated Gene Catalogue^5^, a collection of 9.9 million genes from 1,267 gut metagenome assemblies, which we grouped into 7,063,981 protein clusters at 90% protein identity (referred to as IGC-90). Nearly all samples used to generate the IGC were also included in the UHGP catalogue (except for 59 transcriptome datasets), but the latter was generated from a larger and more geographically diverse metagenomic dataset (including samples from Africa, South America and Oceania). The UHGP-90 and IGC-90 resulted in a combined set of 15.2 million protein clusters, with an overlap of 5.8 million sequences (Fig. 4b). This revealed that 81% of the IGC is represented in the UHGP catalogue, with the missing 19% likely representing fragments of prokaryotic genomes <50% complete, viral or eukaryotic sequences, plasmids or other sequences not binned into MAGs. Most notably though, the UHGP provided an increase of 115% coverage of the gut microbiome protein space over the IGC. As the UHGP was generated from individual genomes and not from their original unbinned metagenome assemblies, our catalogue also has the advantage of providing a direct link between each gene cluster and its genome of origin. This ultimately allows combining individual genes with their genomic context for an integrated study of the gut microbiome.

### Functional capacity of the human gut microbiota

We used the eggNOG^31^, InterPro^32^, COG^33^ and KEGG^34^ annotation schemes to capture the full breadth of functions within the UHGP. However, we found that 42.6% of the UHGP-100 was poorly characterized, as 28.1% lacked a match to any database and a further 14.5% only had a match to a COG with no known function (Fig. 4c). Based on the distribution of COG functions, the most highly represented categories were related to amino acid transport and metabolism, cell wall/membrane/envelope biogenesis and transcription.

We further leveraged the set of 625 million proteins derived from the human gut genomes to explore the functional diversity within each of the UHGG species. Protein sequences from all conspecific genomes were clustered at a 90% amino acid identity to generate a pan-genome for each species. Analysis of the functional capacity of the UHGG species pan-genomes identified a total of 363 KEGG modules encoded by at least one species (Supplementary Fig. 7a and Supplementary Table 4). Most conserved modules were related to ribosomal structure, glycolysis, inosine monophosphate biosynthesis, gluconeogenesis, and the shikimate pathway — all representing essential bacterial functions. However, we found that for certain phyla such as Myxococcota, Bdellovibrionota, Thermoplasmatota, Patescibacteria and Verrucomicrobiota, a substantial proportion of the species pan-genomes remained poorly characterized (Supplementary Fig. 7b). At the same time, species belonging to the clades Fibrobacterota, Bacteroidota, Firmicutes I, Verrucomicrobiota and Patescibacteria had the highest proportion of genes encoding carbohydrate-active enzymes (CAZy; Supplementary Fig. 7b). As most of these lineages are largely represented by uncultured species (Fig. 3b), this suggests the gut microbiota may harbour many species with important metabolic activities yet to be cultured and functionally characterized under laboratory conditions.

### Patterns of intra-species genomic diversity

With the protein annotations and pan-genomes inferred for each of the UHGG species, we explored their intra-species core and accessory gene repertoire. Only near-complete genomes (≥90% completeness) and species with at least 10 independent conspecific genomes were analysed. The overall pattern of gene frequency within each of the 781 species here considered showed a distinctive bimodal distribution (Supplementary Fig. 8), with most genes classified as either core or rare (i.e. present in ≥90% or <10% of conspecific genomes). We analysed the pan-genome size per species in relation to the number of conspecific genomes to look for differences in intra-species gene richness. We observed distinct patterns across different gut phyla, with species from various Firmicutes clades showing the highest rates of gene gain (Fig. 5a). There was a wide variation in the proportion of core genes between species even among clades with more than 1,000 genomes (Fig. 5b), with a median core genome proportion (percentage of core genes out of all genes in the representative genome) estimated at 66% (IQR = 59.6–73.9%).

**Figure 5.**
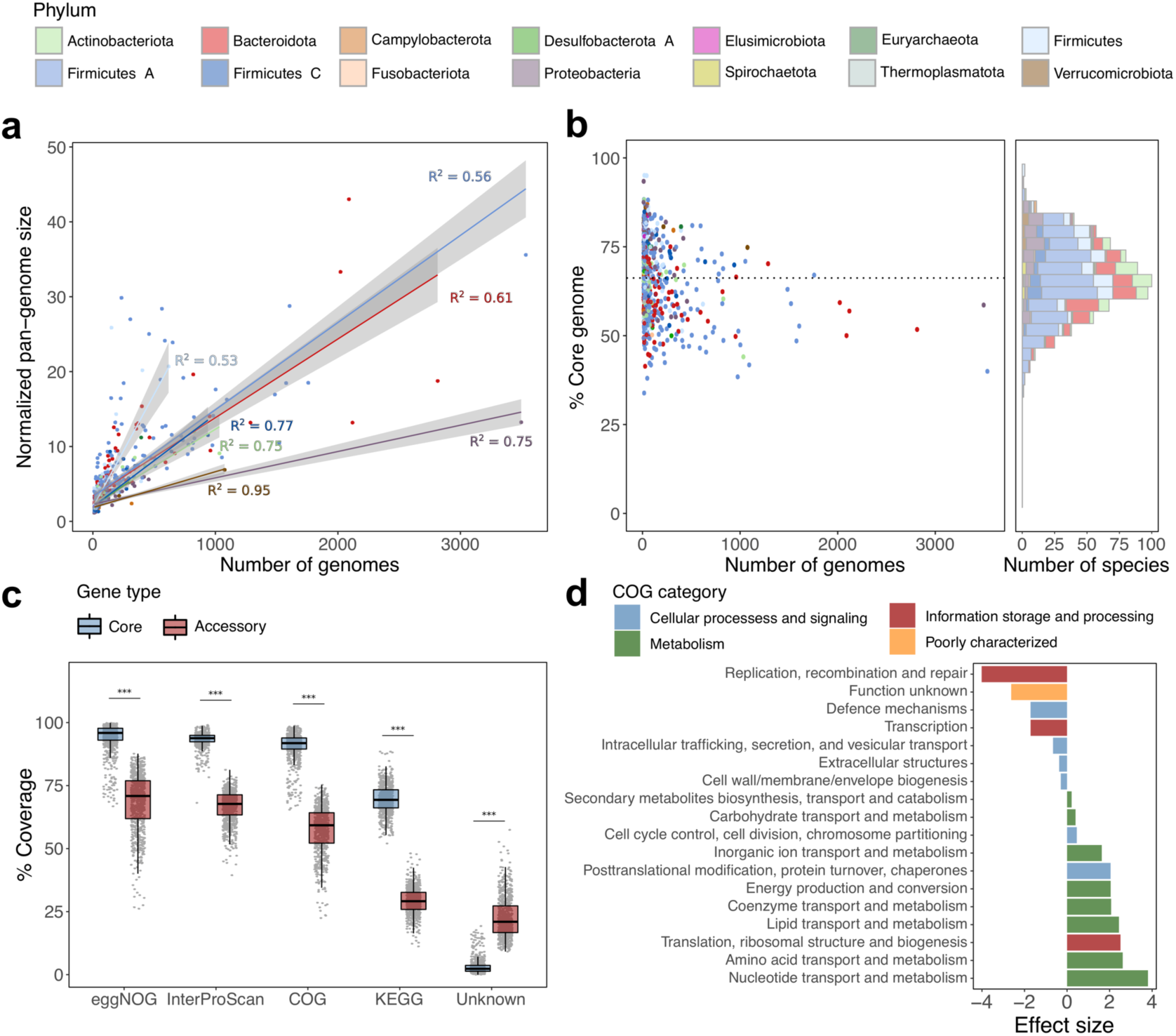
Pan-genome diversity patterns within the gut microbiome. **a**, Normalized pan-genome size as a function of the number of conspecific genomes. Regression curves were generated per phylum, with the corresponding coefficients of determination indicated next to each curve. **b**, Fraction of each species core genome (proportion of core genes out of all genes in the representative genome) according to the number of conspecific genomes (left) and as a histogram (right), coloured by phylum. Horizontal dashed line represents the median value across all species. **c**, Proportion of core and accessory genes from each species that was classified with various annotation schemes, alongside the percentage of genes lacking any functional annotation. \*\*\**P* <0.001 **d**, Comparison between the functional categories assigned to the core and accessory genes. Only those statistically significant (adjusted *P* <0.05) are shown. A positive effect size indicates overrepresentation in the core genes.

To distinguish the functions encoded in the core and accessory genes, we analysed their associated annotations. Core genes were well covered, with a median of 96%, 94%, 92% and 69% of the genes assigned with an eggNOG, InterPro, COG and KEGG annotation, respectively (Fig. 5c). However, the accessory genes had a significantly higher proportion of unknown functions (*P* <0.001), with a median of 21% of the genes (IQR = 16.7–27.3%) lacking a match in any of the databases considered. Thereafter, we investigated the functions encoded by the core and accessory genes on the basis of the COG functional categories. Genes classified as core were significantly associated (adjusted *P* <0.001) with key metabolic functions involved in nucleotide, amino acid and lipid metabolism, as well as other housekeeping functions (e.g. related to translation and ribosomal structure, Fig. 5d). In contrast, accessory genes had a much greater proportion of COGs without a known function, and of genes involved in replication and recombination which are typically found in mobile genetic elements (MGEs, Fig. 5d). A significant number of accessory genes were related to defence mechanisms, which encompass not only general mechanisms of antimicrobial resistance (AMR) such as ABC transporter efflux pumps, but also targeted systems towards invading MGEs (e.g. CRISPR-Cas and restriction modification systems against bacteriophages). These results highlight the potential of this resource to better understand the dynamics of chromosomally encoded AMR within the gut and decipher to what extent the microbiome may be a source of both known and novel resistance mechanisms.

We next investigated intra-species single nucleotide variants (SNVs) within the UHGG species. We generated a catalogue consisting of 249,435,699 SNVs from 2,489 species with three or more conspecific genomes (Fig. 6a). For context, a previously published catalogue contained 10.3 million single nucleotide polymorphisms from 101 gut microbiome species^35^. Of note, more than 85% of these SNVs were exclusively detected in MAGs, whereas only 2.2% were exclusive to isolate genomes (Fig. 6b). We found the overall pairwise SNV density between MAGs to be higher than that observed between isolate genomes (Fig. 6c). Next, we assigned the detected SNVs to the continent of origin of each genome and observed that 36% of the SNVs were continent exclusive. Notably, genomes with a European origin contributed to the most exclusive SNVs (Fig. 6d). However, genomes from Africa contributed over three times more variation on average than European or North American genomes. Pairwise SNV analysis also supported a higher cross-continent SNV density, especially between genomes from Africa and Europe (Fig. 6e). Our results suggest there is a high strain variability between continents and that a considerable level of diversity remains to be discovered, especially from underrepresented regions such as Africa, South America and Oceania.

**Figure 6.**
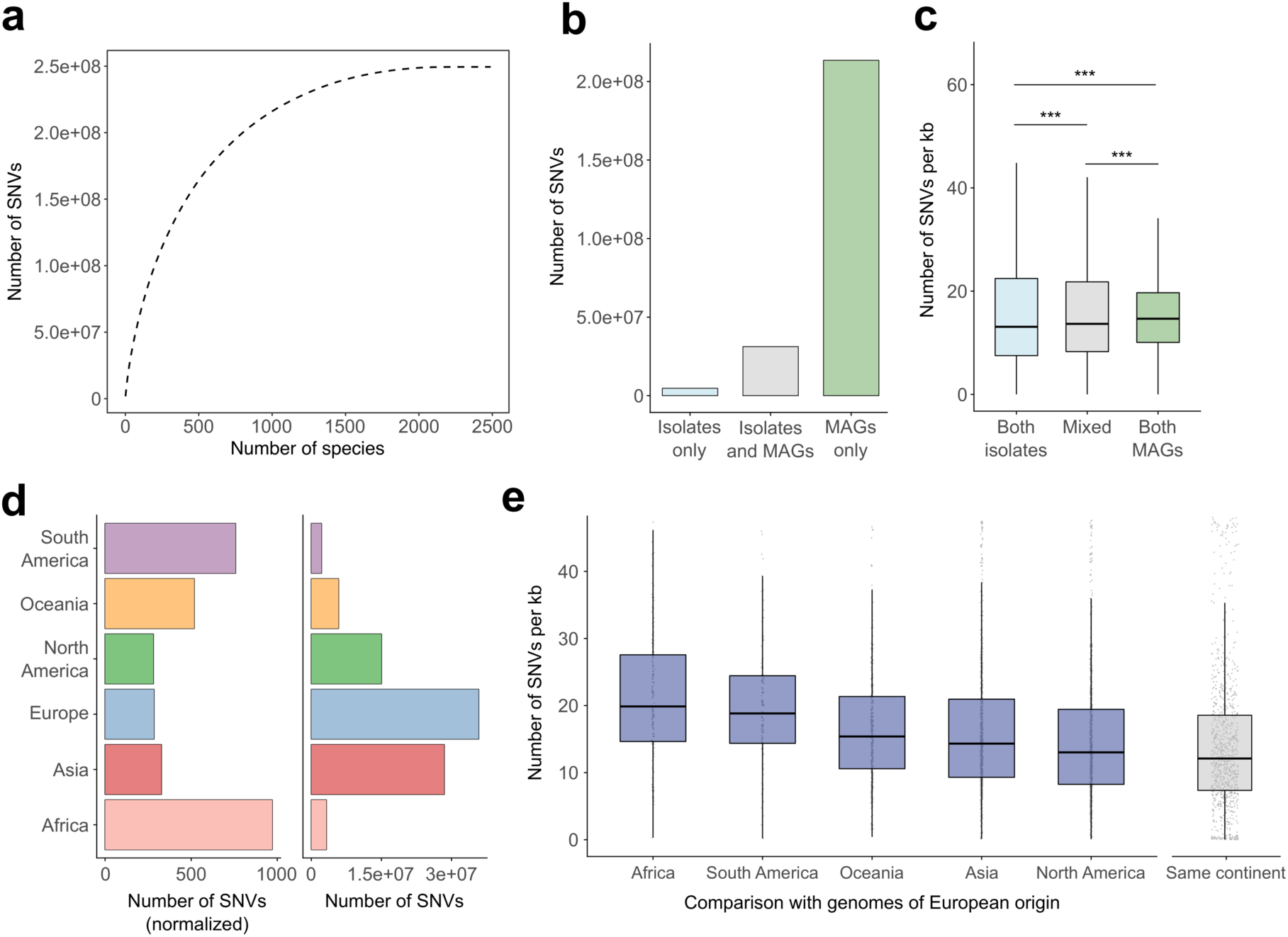
Analysis of intra-species single nucleotide variation. **a**, Total number of SNVs detected as a function of the number of species. The cumulative distribution was calculated after ordering the species by decreasing number of SNVs. **b**, Number of SNVs detected only in isolate genomes, MAGs, or in both. **c**, Pairwise SNV density analysis of genomes of the same or different type. \*\*\**P* < 0.001 **d**, Right panel shows the number of SNVs exclusively detected in genomes from each continent. The left panel shows the number of exclusive SNVs normalized by the number of genomes per continent. **e**, Pairwise SNV density analysis between genomes from Europe, the largest genome subset, and other continents. The median SNV density was calculated per species and the distribution is shown for all species. Comparison of genomes recovered from the same continent was used as reference. The SNV density between genomes of the same continent is significantly lower (adjusted *P* < 0.05) to that calculated for genomes from different continents.

### Resource implementation

Both the UHGG and UHGP catalogues are available as part of a new genome layer within the MGnify^36^ website, where summary statistics of each species cluster and their functional annotations can be interactively explored and downloaded (see ‘Data availability’ section for more details). We have also generated a BItsliced Genomic Signature Index (BIGSI)^37^ of the UHGG, which will allow users to interactively screen for the presence of small sequence fragments (<5 kb) in this collection. As new genomes from the human gut microbiome are generated and made publicly available, we plan to periodically update the resource with newly discovered species or by replacing uncultured reference genomes with better quality versions.

## Discussion

We have generated a unified sequence catalogue representing 286,997 genomes and over 625 million protein sequences of the human gut microbiome. Of the 4,644 species contained in the UHGG, 71% lack a cultured representative, meaning the majority of microbial diversity in the catalogue remains to be experimentally characterized. During preparation of our manuscript, a new collection of almost 4,000 cultured genomes from 106 gut species was released^38^, which will be incorporated in future versions of the resource. As 96% of these genomes were reported to have a species representative in the culture collections here included, we do not anticipate this dataset to provide a substantial increase in the number of species discovered. Nevertheless, our analyses suggest additional uncultured species from the human gut microbiome are yet to be discovered, highlighting the importance and need for culture-based studies. Furthermore, given the sampling bias towards populations from China, Europe and the United States, we expect that many underrepresented regions still contain substantial uncultured diversity.

By comparing recently published large datasets of uncultured genomes^16,17,19^, we were able to assess the reproducibility of the results from each study. We show that despite the different assembly, binning and refinement procedures employed in the three studies, almost all of the same species and near-identical strains were recovered independently when using a consistent sample set. These results further increase confidence in the use of metagenome-assembled genomes for the characterization of uncultured microbial diversity.

With the establishment of this massive sequence catalogue, it is evident that a large portion of the species and functional diversity within the human gut microbiome remains uncharacterized. Moreover, our knowledge of the intra-species diversity of many species is still limited due to the presence of a small number of conspecific genomes. Having this combined resource can help guide future studies and prioritize targets for further experimental validation. Using the UHGG or UHGP, the community can now screen for the prevalence and abundance of species/genes in a large panel of intestinal samples and in specific clinical contexts. By pinpointing particular taxonomic groups with biomedical relevance, more targeted approaches could be developed to improve our understanding of their role in the human gut. The functional predictions generated for the species pan-genomes could also be leveraged to develop new culturing strategies for isolation of candidate species. Target-enrichment methods such as single-cell^39^ and/or bait-capture hybridization^40^ approaches could also be applied. Being able to enrich for specific groups of interest, even without culturing, could allow recovery of better-quality versions of MAGs and improve the analysis derived from genome sequence data alone. Given the large uncultured diversity still remaining in the human gut microbiome, having a high-quality catalogue of all currently known species substantially enhances the resolution and accuracy of metagenome-based studies. Therefore, the presented genome and protein catalogue represents a key step towards a hypothesis-driven, mechanistic understanding of the human gut microbiome.

## Methods

### Genome collection

We compiled all the prokaryotic genomes publicly available as of March 2019 that have been sampled from the human gut. To retrieve isolate genomes, we surveyed the IMG^23^, NCBI^21^ and PATRIC^22^ databases for genome sequences annotated as having been isolated from the human gastrointestinal tract. We complemented this set with bacterial genomes belonging to two recent culturomics collections: the Human Gastrointestinal Bacteria Culture Collection (HBC)^18^ and the Culturable Genome Reference (CGR)^20^. To avoid including duplicated entries due to redundancy between reference databases, we combined genomes obtained from the PATRIC and IMG repositories, and added only those without an identical genome in the sets extracted from NCBI, HBC and CGR. Metagenome-assembled genomes (MAGs, i.e. uncultured genomes) were obtained from Pasolli, et al.^19^ (CIBIO), Almeida, et al.^17^ (EBI) and Nayfach, et al.^16^ (HGM). For the CIBIO set, only those genomes retrieved from samples collected from the intestinal tract were used. Metadata for each genome was retrieved using the API of the various public repositories and combined with that available in each of the original studies.

### Assessing genome quality

Genome quality (completeness and contamination) was estimated with CheckM v1.0.11^41^ using the ‘lineage_wf’ workflow to select only those that passed the following criteria: >50% genome completeness, <5% contamination and an estimated quality score (completeness – 5 × contamination) >50. We also searched for the presence of ribosomal RNAs in each genome with the ‘cmsearch’ function of INFERNAL^42^ (options ‘-Z 1000 --hmmonly --cut_ga --noali – tblout’) against the Rfam^43^ covariance models for the 5S, 16S and 23S rRNAs. tRNAs of the standard 20 amino acids were identified with tRNAScan-SE^44^ with options ‘-A -Q’ for archaeal species and ‘-B -Q’ for those belonging to bacterial lineages.

### Species clustering

We clustered the total set of 286,997 genomes at an estimated species level (average nucleotide identity, ANI ≥95%^25^) using dRep v2.2.4^45^ with the following options: ‘-pa 0.9 -sa 0.95 -nc 0.30 -cm larger’. Because of the computational burden of clustering together the entire genome set, we employed an iterative approach where random chunks of 50,000 genomes were clustered independently. The selected representatives from each chunk were combined and subsequently clustered, reducing the final computational load. To ensure the best quality genome was selected as the species representative in each iteration, a score was calculated for each genome based on the following formula:

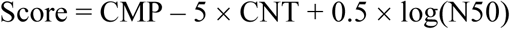

where CMP represents the completeness level, CNT the estimated contamination and N50 the assembly contiguity characterized by the minimum contig size in which half of the total genome sequence is contained. The genome with the highest score was chosen as the species representative, with cultured genomes prioritized over uncultured genomes (i.e. if a MAG had a higher score than an isolate genome, the latter would still be chosen as the representative).

### Evaluating methods reproducibility

The species clusters inferred here were compared with those previously generated in the human gut MAG studies^16,17,19^ from a common set of genomes. Similarity between species clusterings was estimated using the Adjusted Rand Index (ARI) computed in the Scikit-learn python package^46^. This metric considers both the number of clusters and cluster membership to compute a similarity score ranging from 0 to 1.

Conspecific genomes recovered in the same metagenomic samples but in different studies were compared with FastANI v1.1^25^ with default parameters to obtain both the maximum aligned fraction and ANI for each pairwise comparison.

### Inferring cultured status

To determine their cultured status, the UHGG species representatives were searched against NCBI RefSeq release 93 after excluding uncultured genomes (i.e. metagenome-assembled or single-cell amplified genomes). Genome alignments were performed in two stages: (1) Mash v2.1 was used as an initial screen (using the function ‘mash dist’) to identify the most similar RefSeq genome to each of the UHGG species, and (2) ‘dnadiff’ from MUMmer v4.0.0beta2^47^ was subsequently used to compute whole genome ANI between the genome pairs. A species was considered to have been cultured if (1) it contained a cultured gut genome from the UHGG catalogue, or (2) if it matched an isolate RefSeq genome with at least 95% ANI over at least 30% of the genome length.

### Calculating number of conspecific genomes

For an accurate assessment of the number of non-redundant genomes belonging to each species, we de-replicated all conspecific genomes at a 99.9% ANI threshold using dRep with options ‘-pa 0.999 –SkipSecondary’. Furthermore, the frequency of each species was only counted once per sample to avoid cases where the same genome was recovered multiple times because of overlapping samples between the three MAG studies.

### Estimating geographical diversity

A geographical diversity index was estimated to assess how widely distributed each species was. We calculated the Shannon diversity index on the proportion of samples each species was found per continent. This metric combines both richness and evenness, so the level of estimated diversity is highest in species found across all continents at a similar proportion.

### Phylogenetic analyses

Taxonomic annotation of each species representative was performed with the Genome Taxonomy Database^27^ Toolkit (GTDB-Tk) v0.3.1 (https://github.com/Ecogenomics/GTDBTk) (database release 04-RS89) using the ‘classify_wf’ function and default parameters. To use consistent species boundaries between the genome clustering and taxonomic classification procedures, genomes were assigned at the species level if the ANI to the closest GTDB-Tk species representative genome was ≥95% and the alignment fraction ≥30%. In this taxonomy scheme, genera and species names with an alphabetic suffix indicate taxa that are polyphyletic or needed to be subdivided based on taxonomic rank normalization according to the current GTDB reference tree. The lineage containing the type strain retains the unsuffixed (valid) name and all other lineages are given alphabetic suffixes, indicating they are placeholder names that need to be replaced in due course. Taxon names above the rank of genus appended with an alphabetic suffix indicate groups that are not monophyletic in the GTDB reference tree, but for which there exists alternative evidence that they are monophyletic groups. We also generated NCBI taxonomy annotations for each species-level genome based on its placement in the GTDB tree, using the ‘gtdb_to_ncbi_majority_vote.py’ script available in the GTDB-Tk repository (https://github.com/Ecogenomics/GTDBTk/tree/stable/scripts).

Maximum-likelihood trees were generated *de novo* using the protein sequence alignments produced by the GTDB-Tk: we used IQ-TREE v1.6.11 to build a phylogenetic tree of the 4,616 bacterial and 28 archaeal species. The best-fit model was automatically selected by ‘ModelFinder’ on the basis of the Bayesian Information Criterion (BIC) score. The LG+F+R10 model was chosen for building the bacterial tree, while the LG+F+R4 model was used for the archaeal phylogeny. Trees were visualized and annotated with the Interactive Tree Of Life (iTOL) v4.4.2^48^. Phylogenetic diversity (PD) was estimated by the sum of branch lengths, with the amount that was exclusive to uncultured species calculated as PD_total_ – PD_cultured_. Uncultured monophyletic groups were defined as nodes in the tree containing child leaves exclusively comprised of uncultured genomes.

### BIGSI construction

A BItsliced Genomic Signature Index (BIGSI)^37^ was generated for all species-level genomes with BIGSI v0.3.8. First, *k*-mers of size 31 were extracted from each genome with McCortex v1.0.1^49^ (‘mccortex31 build -k 31’). Thereafter, Bloom filters were built for each *k*-mer set using ‘bigsi bloom’ and inserted into the BIGSI index with ‘bigsi build’. BIGSI config parameters *h* (number of hash functions applied to each *k*-mer) and *m* (Bloom filter’s length in bits) were set at 1 and 28,000,000, respectively. A final API layer for querying the index was built using hug (http://www.hug.rest/) and hosted on the MGnify^36^ website: https://www.ebi.ac.uk/metagenomics/genomes.

### Pan-genome analysis and functional annotation

Protein coding sequences (CDS) for each of the 286,997 genomes were predicted and annotated with Prokka v1.13.3^50^, using Prodigal v2.6.3^51^ with options ‘-c’ (predict proteins with closed ends only), ‘-m’ (prevent genes from being built across stretches of sequences marked as Ns) and ‘-p single’ (single mode for genome assemblies containing one single species). Pan-genome analyses were carried out using Roary v3.12.0^52^. We set a minimum amino acid identity for a positive match at 90% (‘-i 90’), a core gene defined at 90% presence (‘-cd 90’) and no paralog splitting (‘-s’). A normalized pan-genome size was estimated by dividing the total number of core and accessory genes by the number of genes contained in the species representative genome.

The Unified Human Gastrointestinal Protein (UHGP) catalogue was generated from the combined set of 625,251,941 CDS predicted. Protein clustering of the UHGP and the Integrated Gene Catalogue (IGC)^5^ was performed with the ‘linclust’ function of MMseqs2 v6-f5a1c^53^ with options: ‘--cov-mode 1 -c 0.8’ (minimum coverage threshold of 80% length of the shortest sequence) and ‘--kmer-per-seq 80’ (number of *k*-mers selected per sequence, increased from the default of 21 to improve clustering sensitivity). The ‘--min-seq-id’ option was set at 1, 0.95, 0.9 and 0.5 to generate the catalogues at 100%, 95%, 90% and 50% protein identity, respectively. We clustered the IGC solely at a 90% and 50% protein identity as it was originally de-replicated at a 95% nucleotide identity^5^. Functional characterization of all protein sequences was performed with eggNOG-mapper v2^54^ (database v5.0^31^) and InterProScan v5.35-74.0^32^. COG^33^, KEGG^34^ and CAZy^55^ annotations were derived from the eggNOG-mapper results. Differences in annotation coverage and COG functional categories between the core and accessory genes were evaluated with a two-tailed Wilcoxon rank-sum test in R v3.6.0 (function ‘wilcox.test’). Expected *P* values were corrected for multiple testing with the Benjamini-Hochberg method. Cohen’s d effect sizes were estimated with the function ‘cohen.d’ from the Effsize^56^ R package. To accurately estimate the proportion of each KEGG module in the species pan-genome, we used the compositional data analysis R package CoDaSeq^57^. Pseudo counts for zero-count data were first imputed using a Bayesian-Multiplicative simple replacement procedure implemented in the ‘cmultRepl’ function (method ‘CZM’). Final counts were thereby converted to centred log-ratios using the ‘codaSeq.clr’ function to account for the compositional nature of the data and for differences in pan-genome size.

### SNV analyses

A total of 2,489 species with at least three conspecific genomes were used for generating a catalogue of single nucleotide variants (SNVs). For each species, we mapped all conspecific genomes to the representative genome using the ‘nucmer’ program from MUMmer v4.0.0.beta2^47^ and filtered alignments using the ‘delta-filter’ program with options ‘-q -r’ to exclude chance- and repeat-induced alignments. Thereafter, we identified SNVs using the ‘show-snps’ program. Single base insertions and deletions were not counted as SNVs. Each SNV locus was included in the catalogue only when the alternate allele was detected in at least two conspecific genomes. The final SNV catalogue was generated by unifying the SNV coordinates on the basis of their position in the species representative genome. The SNV entries in the catalogue were characterized as genome type-specific or continent-specific based on whether the alternate allele could be found solely in genomes from a specific genome type or continent. The number of continent-specific SNVs was normalized by the number of genomes from the corresponding continent to estimate the contribution per genome to the continent-specific SNV discoveries.

Similar programs and parameters were used for the pairwise genome alignment, but in this case only near-complete genomes (≥90% completeness) and species with at least 10 independent conspecific genomes were considered. Due to the high computational demand, pairwise alignments of species encompassing more than 1,000 genomes were limited to the best-quality 1,000 genomes. A total of 29,283,684 pairwise genome alignments were performed between almost 113,000 genomes from 909 species. For each pairwise comparison, we estimated the total number of SNVs and the overall density as the number of SNVs per kb. In addition, the pairwise comparisons were organized based on the type and the continent origin of the genomes in the pair for further downstream analyses. A two-tailed Wilcoxon rank-sum test was used to evaluate differences in SNV distributions. Resulting *P* values were corrected for multiple testing with the Benjamini-Hochberg method.

### Data availability

Genome assemblies of the UHGG have been deposited in the European Nucleotide Archive under study accession ERP116715. The UHGG, UHGP and SNV catalogues are available in a public FTP server (http://ftp.ebi.ac.uk/pub/databases/metagenomics/mgnify_genomes/) alongside functional annotations and the pan-genome results. These data together with the BIGSI search index of the UHGG can also be accessed interactively on the MGnify website: https://www.ebi.ac.uk/metagenomics/genomes.

## Acknowledgements

We would like to thank Phelim Bradley and Zamin Iqbal for their help in the BIGSI implementation, and Dongying Wu for assistance in the identification of uncultured monophyletic groups. Funding: European Molecular Biology Laboratory (EMBL); Biotechnology and Biological Sciences Research Council [BB/N018354/1 and BB/R015228/1]; European Research Council (project ERC-STG MetaPG-716575) to N.S.

## Author contributions

A.A., S.N., N.K.C. and R.D.F. conceived the study. A.A. performed the genome clustering and annotations, compared study sets, carried out the pan-genome analyses, built the BIGSI index and drafted the manuscript. S.N. provided feedback, performed phylogenetic, rarefaction and clustering analyses, as well as the comparison with RefSeq. M.Boland and M.Beracochea built the resource implementation within the MGnify website. F.S. built the protein catalogue and performed the comparison with the IGC. Z.J.S. generated the SNV catalogue and performed related analyses according to genome type and geographic origin. K.S.P. provided feedback, funding and contributed to the SNV analyses. D.H.P. and P.H. provided feedback and assisted in the species taxonomic classification. N.S. provided feedback, funding and contributed to the generation of the protein catalogue. N.K.C. and R.D.F. supervised the work, provided feedback and funding. All authors read, edited and approved the final manuscript.

## Competing interests

F.S. is an employee of Enterome SA. P.H. is a co-founder and Director of Microba Life Sciences Limited. D.H.P. is a consultant to Microba Life Sciences Limited. R.D.F. is a consultant to Microbiotica Pty Ltd.

## Supplementary Figures

**Supplementary Figure 1.**
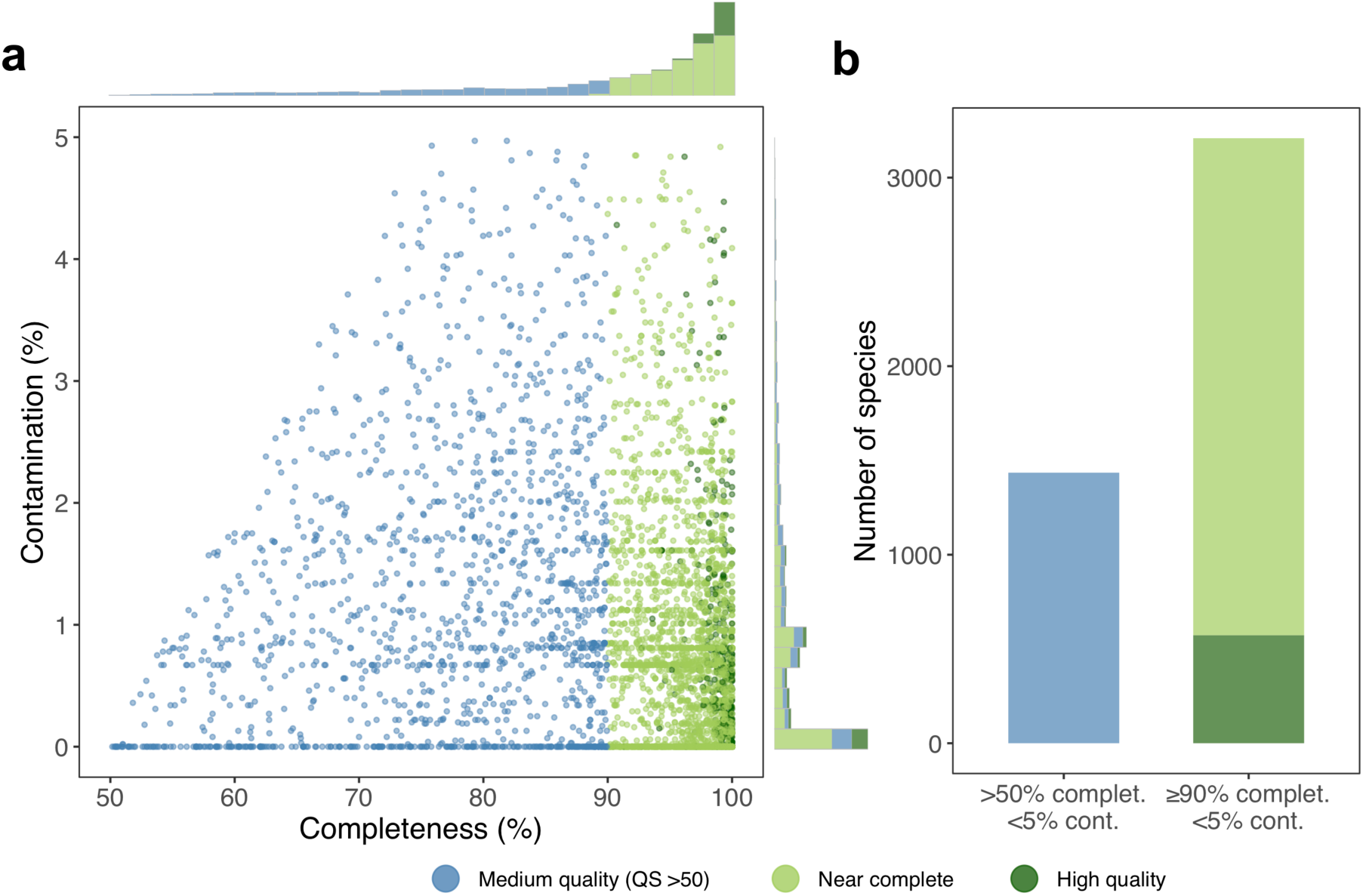
Genome quality of species representatives. **a**, Completeness and contamination scores for each of the 4,644 species representatives, coloured by their quality classification category. Medium quality: >50% completeness; near complete: ≥90% completeness; high-quality: >90% completeness, presence of 5S, 16S and 23S rRNA genes, as well as at least 18 tRNAs. All genomes have a quality score (QS = completeness – 5 × contamination) above 50. **b**, Number of species according to different completeness and contamination criteria.

**Supplementary Figure 2.**
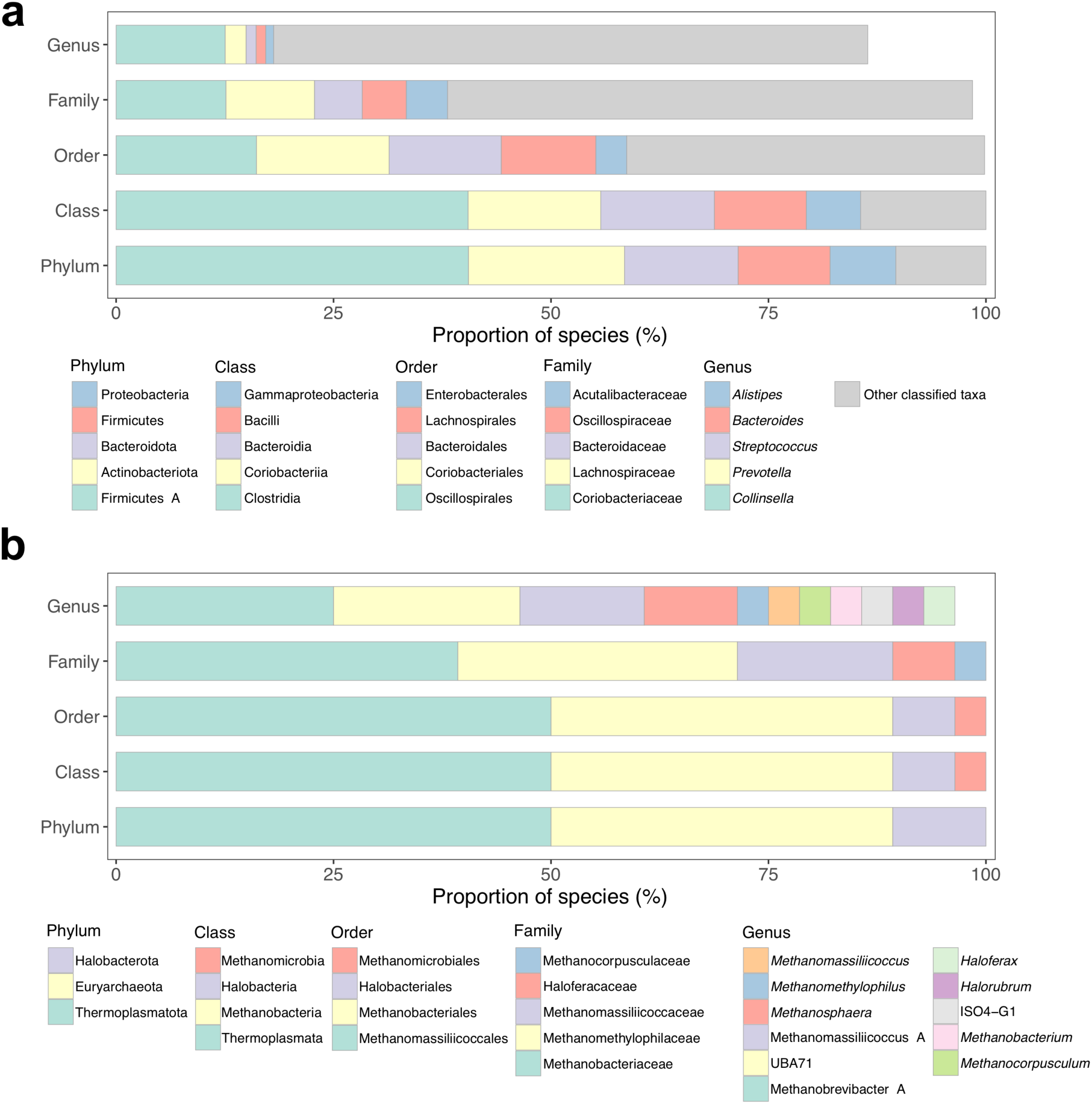
Taxonomy composition of the bacterial and archaeal species. **a**, Taxonomic affiliation of the 4,616 bacterial species detected. Data is partitioned by taxonomic rank, with only the five most highly represented taxa per rank depicted in the legend. **b**, Taxonomic affiliation of the 28 archaeal species detected, partitioned by taxonomic rank.

**Supplementary Figure 3.**
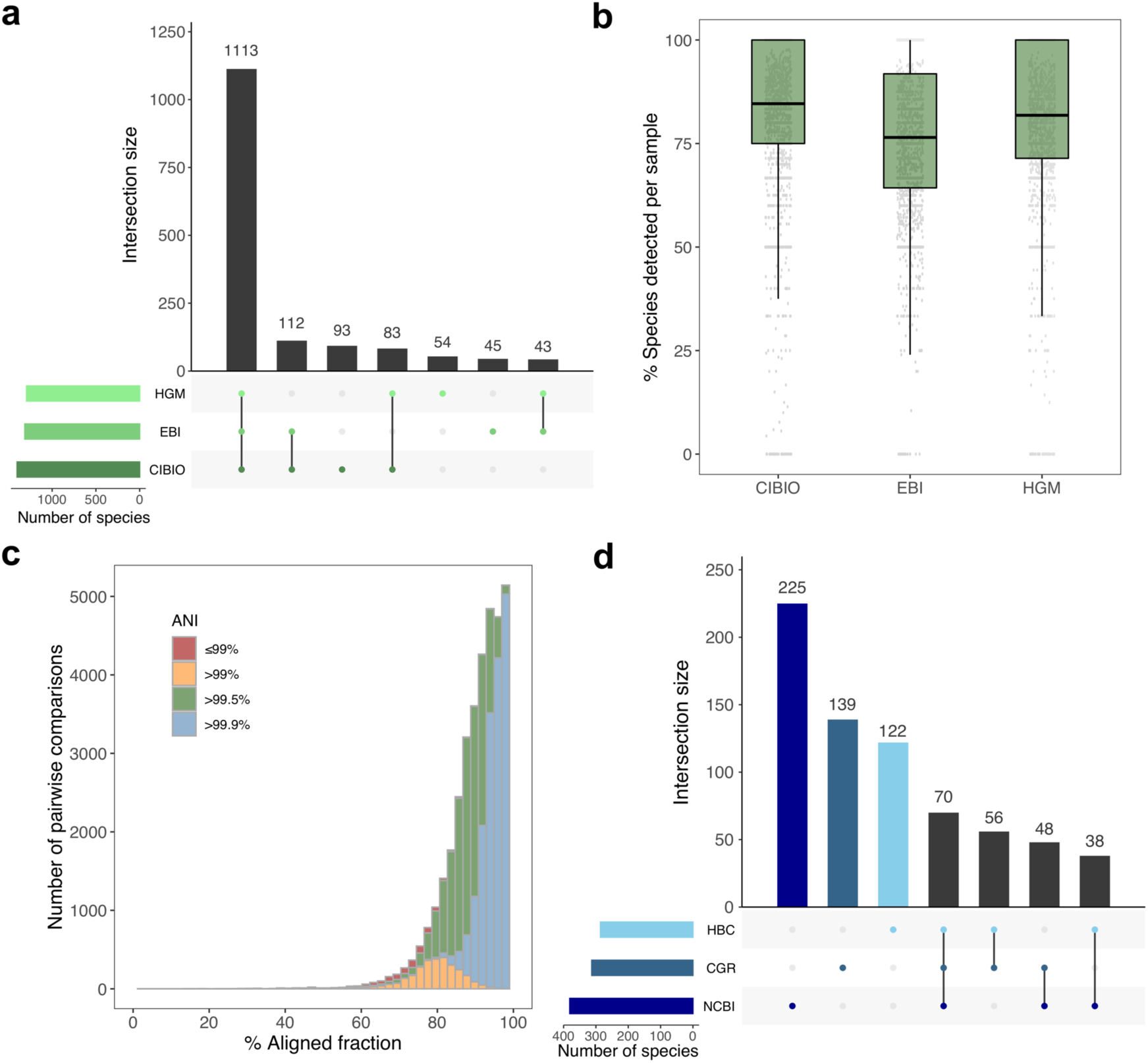
Species overlap across study sets. **a**, Number of species found across the three metagenome-assembled genome sets, ordered by their level of overlap. Only those genomes recovered from the 1,554 metagenomic samples used by all three studies were considered in this analysis. **b**, Distribution of the proportion of species recovered per sample in each study out of all species recovered across all three studies in the same samples. **c**, Estimated aligned fractions and average nucleotide identities (ANI) between conspecific genomes obtained in the same sample but in different MAG studies. **d**, Number of species identified in three culture-based studies and their degree of overlap. The NCBI study set consists mainly of genomes from the Human Microbiome Project (HMP).

**Supplementary Figure 4.**
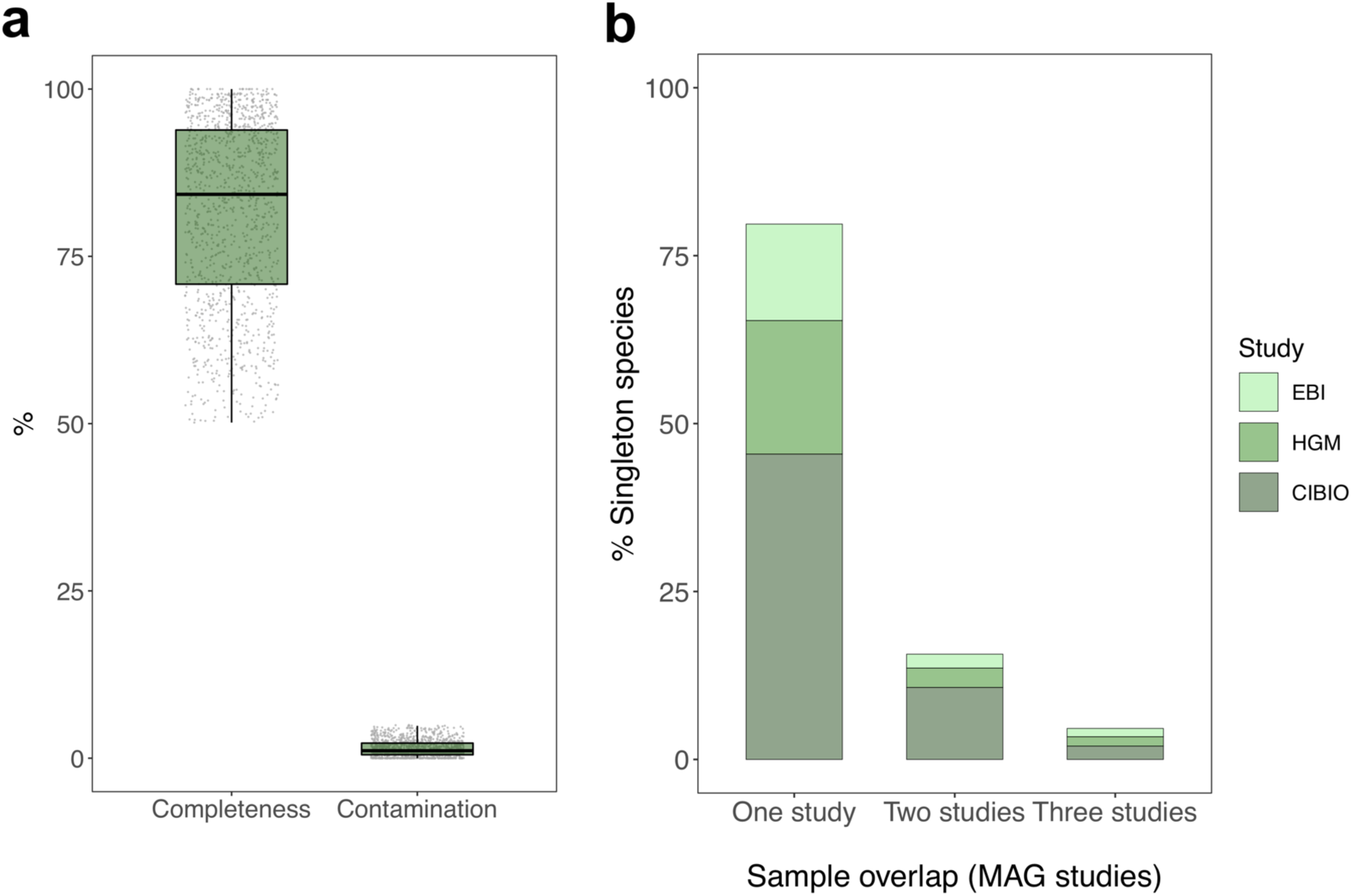
Quality and sample origin of uncultured singleton species. **a**, Genome completeness and contamination estimates of the 1,212 uncultured species represented by a single genome. **b**, Proportion of the 1,212 singleton species, by study set, that originated from samples analysed in one, two or three of the MAG studies (CIBIO, EBI and HGM). The CIBIO study used metaSPAdes and MetaBAT 2 for assembling and binning sequencing runs previously merged by sample; the HGM study used MEGAHIT to assemble runs merged by sample and applied a combination of MaxBin 2, MetaBAT 2, CONCOCT and DAS Tool for binning and refinement; the EBI study used metaSPAdes and MetaBAT 2 for assembling and binning individual runs without merging by sample.

**Supplementary Figure 5.**
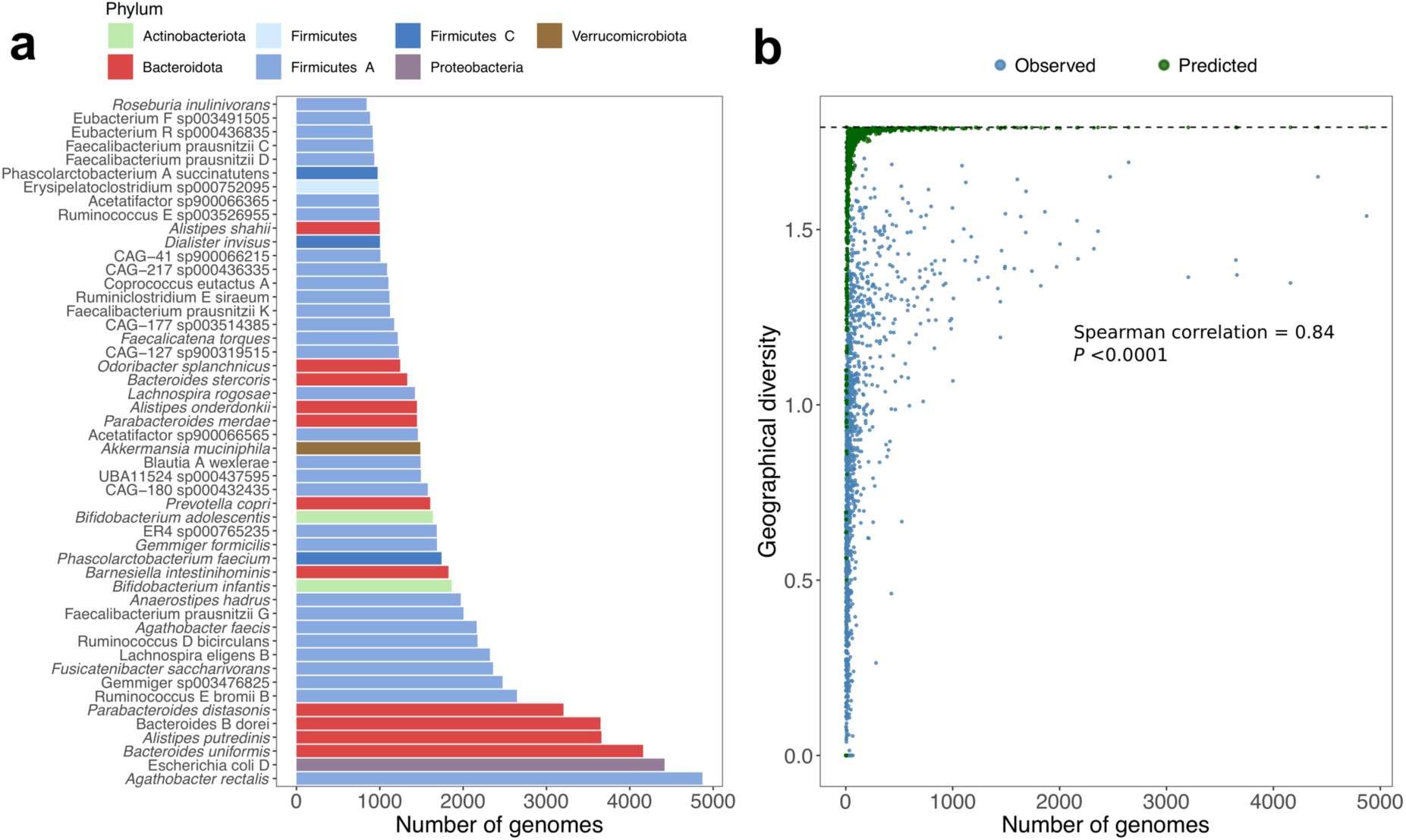
Species frequency and geographical diversity. **a**, Number of non-redundant genomes retrieved from the 50 most highly represented species in the UHGG. Each species is coloured by its assigned phylum according to the figure legend. **b**, Geographical diversity estimated using the Shannon index in relation to the number of non-redundant genomes from each species. The Spearman’s rank correlation coefficient and *P* value are depicted in the graph. Predicted values represent the random geographical distribution of equivalent numbers of genomes observed for each species. Dashed horizontal line indicates the maximum theoretical value of geographical diversity corresponding to equal sample proportions across the six major continents (Africa, Asia, Europe, North America, South America and Oceania).

**Supplementary Figure 6.**
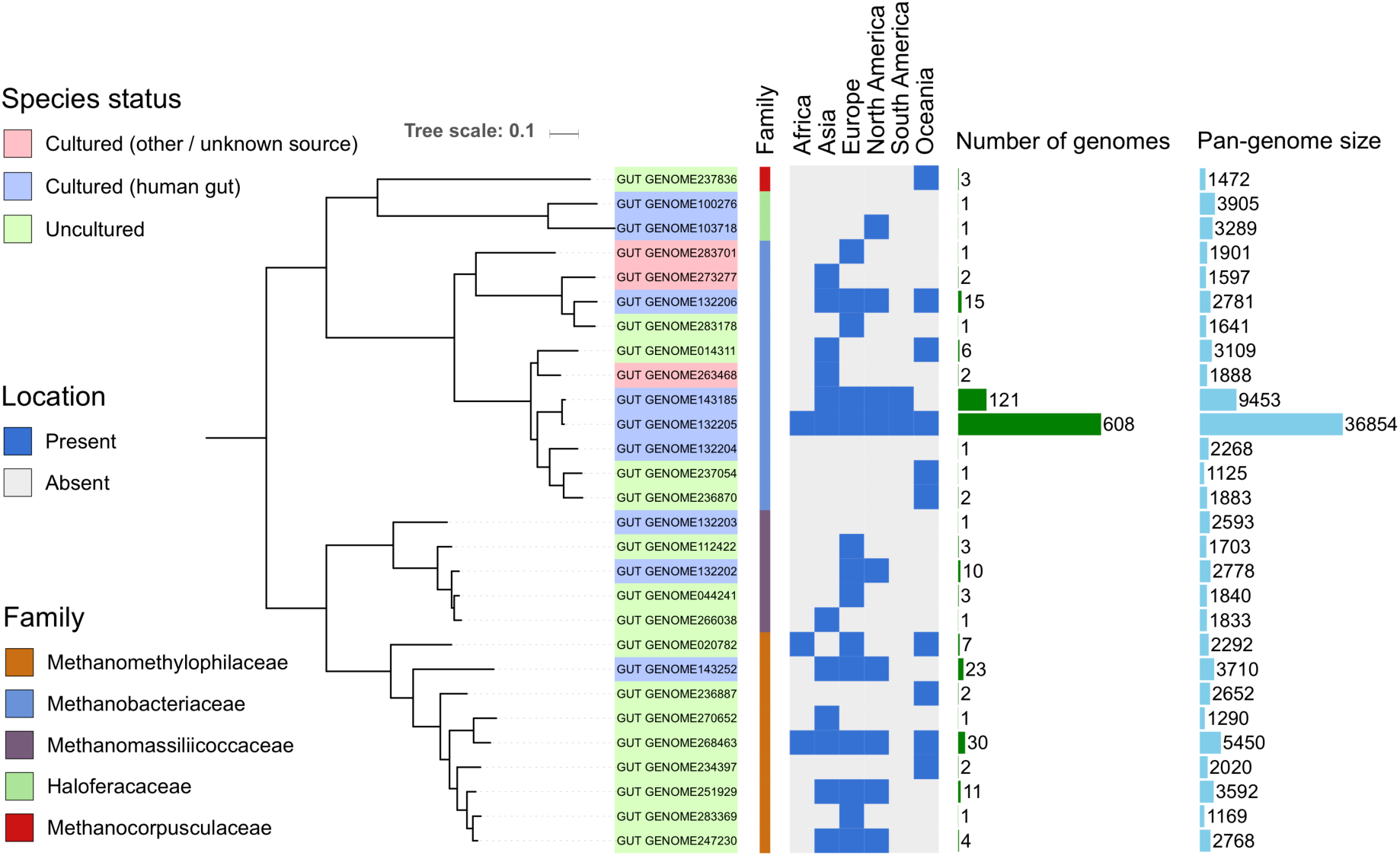
Diversity of the gut archaeal species detected. Phylogenetic tree of the 28 archaeal species detected in the human gut. Tips are labelled with the corresponding species representative code and coloured according to its cultured status. The taxonomic affiliation (family), geographical distribution, number of non-redundant genomes and total pan-genome size are represented next to the tree.

**Supplementary Figure 7.**
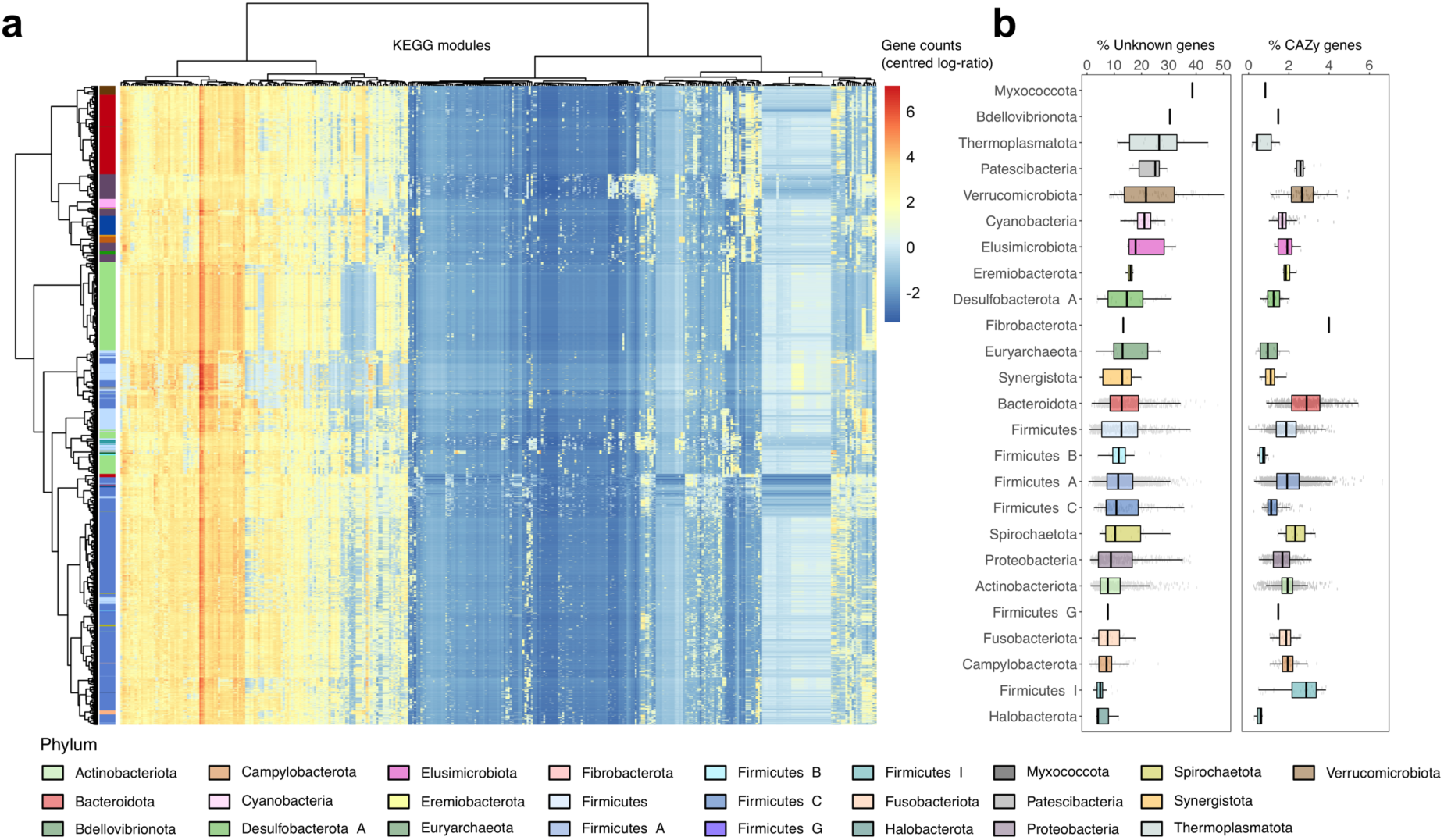
Functional annotation of gut microbiome species. **a**, Functional profiles of the UHGG species pan-genomes (rows) according to 363 KEGG modules (columns). Numbers of genes matching each module were normalized to centred log-ratios after imputing values with zero counts. Species are coloured according to phylum. KEGG modules and species were hierarchically clustered using the Ward’s criterion method. **b**, Proportion of each species pan-genome, partitioned by phylum, without any assignment to the eggNOG, InterPro, COG or KEGG databases (left). Proportion of the pan-genome with a match to the carbohydrate-active enzymes (CAZy) database (right).

**Supplementary Figure 8.**
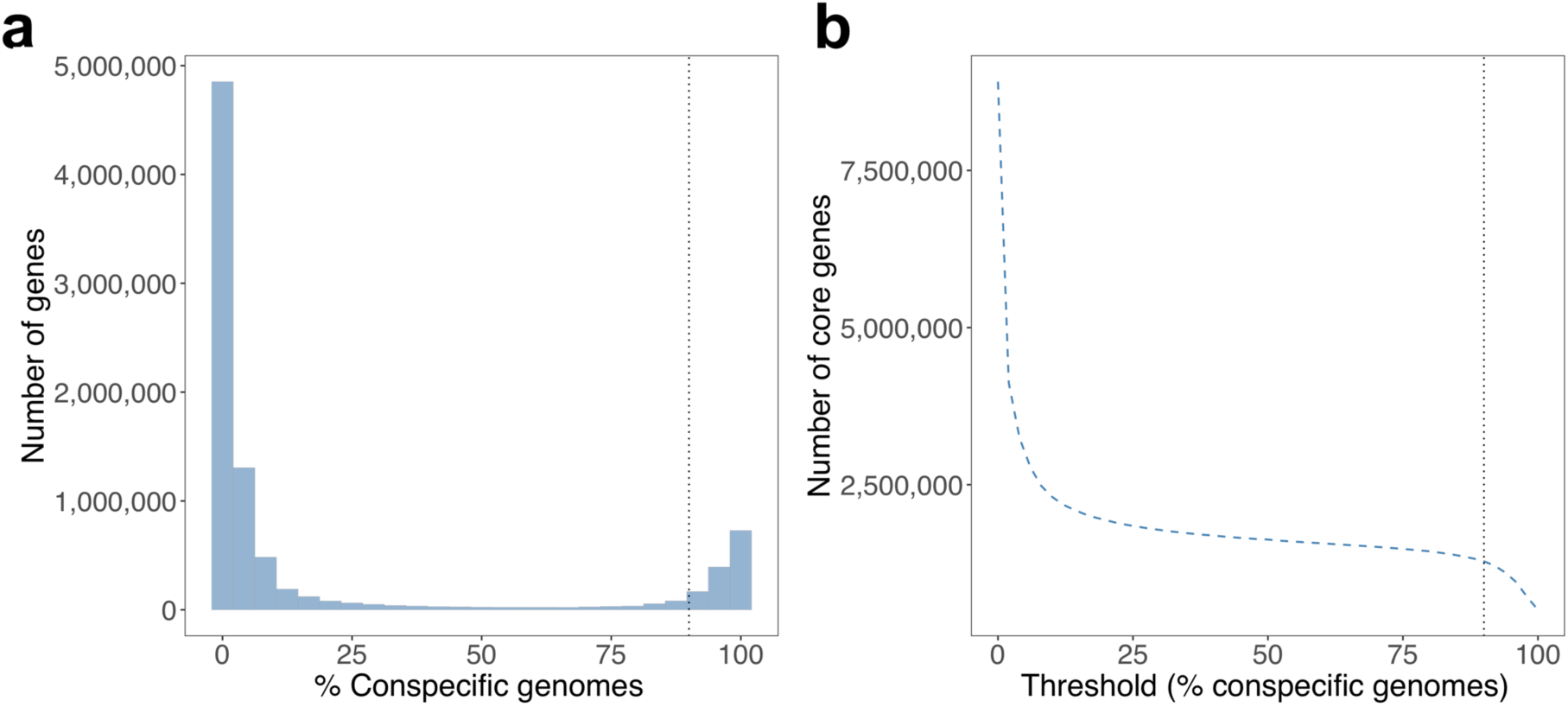
Gene frequency distribution within the species-level clusters. **a**, Distribution of the number of genes found per fraction of conspecific genomes. Only near-complete genomes (≥90% completeness) were considered in the analysis. **b**, Number of core genes detected based on the threshold of genomes per species used to classify as core. Vertical dashed line represents the 90% threshold used in this study.

## Notes

http://ftp.ebi.ac.uk/pub/databases/metagenomics/mgnify_genomes

https://www.ebi.ac.uk/metagenomics/genomes

